# Methods for Analyzing Imaging Mass Cytometry Data with Tellurium Probes

**DOI:** 10.1101/567685

**Authors:** Jay Bassan, Mark Nitz

## Abstract

Imaging mass cytometry (IMC) is a technique allowing visualization and quantification of over 40 biological parameters in a single experiment with subcellular spatial resolution, however most IMC experiments are limited to endpoint analysis with antibodies and DNA stains. Small molecules containing tellurium are promising probes for IMC due to their cell permeability, synthetic versatility, and most importantly their application to sequential labelling with isotopologous probes (SLIP) experiments. SLIP experiments with tellurium-containing probes allow quantification of intracellular biology at multiple timepoints with IMC. Despite the promise of tellurium in IMC, there are unique challenges in image processing associated with tellurium IMC data. Here, we address some of these issues by demonstrating the removal of xenon background signal, combining channels to improve signal-to-noise ratio, and calculating isotope transmission efficiency biases. These developments add accuracy to the unique temporal resolution afforded by tellurium IMC probes.

## Introduction

The emerging technology of imaging mass cytometry (IMC) has delivered insight into many aspects of biology including the heterogeneity of breast cancer tumours and the tissue distribution of cisplatin (1, 2). Tissue sections are stained with more than 40 different antibodies, each of which is conjugated to a polymer that chelates a distinct elemental isotope, typically from the lanthanide series. The section is then ablated pixelwise by a rastering, pulsed ultraviolet laser. The ablated material from each pixel passes into an inductively-coupled plasma mass spectrometer and each isotope, which corresponds to a specific antibody, is quantified. A multi-channel image is thereby created with many more channels accessible than traditional immunohistochemical or immunofluorescent optical imaging.

This technique has been limited to the imaging of static biological markers (i.e. proteins), DNA, or certain small molecule probes against which custom antibodies have been raised (e.g. EF5) (1, 3). With tellurium as an IMC-visible element, we have developed tellurophenes as biologically-compatible mass tags whose imaging requires no antibody staining (4). Probes for specific biological processes can be synthesized by linking tellurophenes, which are aromatic, stable, and non-toxic, to activity-based functional groups that covalently bind cells of a certain phenotype. In this way, IMC can be used to visualize processes or microenvironments including protein synthesis and cellular hypoxia (5, 6). Imaging of these compounds is not dependent on antibody-epitope binding and as such represents direct visualization of the probe itself.

Tellurium exists naturally as a mixture of eight stable isotopes, of which six are commerically available in an isotopically-enriched form. By synthesizing isotopically-pure probes, any given process can be investigated with spatial and temporal resolution with IMC. For example, changes in hypoxia over time were quantified by dosing one isotope of a tellurophene probe conjugated to a 2-nitroimidazole group, followed by waiting or an intervention, followed by a dose of the same molecule with a different isotope (6). The single mass unit resolution of IMC then allows quantification of the differences in labelling of the probes. We term this approach sequential labelling of isotopologous probes (SLIP). SLIP experiments with tellurium add temporal resolution to the IMC toolbox, which is crucial in understanding the deep biological profile imaged by IMC.

While software for analyzing IMC data exist (7, 8), they do not deal explicitly with tellurium as an analyte. Given the specific benefits of tellurophene probes, we sought to design data processing strategies that would improve the accuracy of research in this field and allow SLIP experiments to be included as robust methods in the IMC community.

### Experimental Methods

All appropriate ethical guidelines were followed for animal experiments. Full experimental details describing the preparation of the tissue samples will be described elsewhere. Briefly, mice were injected intravenously with a tellurium-containing probe. After 1–3 hours, animals were sacrificed. The tissues of interest were excised, fixed in formalin, embedded in paraffin, and 5 µm sections were obtained. The formalin-fixed paraffin-embedded tissue sections were prepared for IMC as per Fluidigm’s suggested protocols (Protocol ID PN 400322 A3). Stained tissue sections were imaged on the Hyperion^TM^ Imaging System at 200 Hz. The software controlling the instrument outputs a .txt file which was then processed as described in this manuscript.

## Results and Discussion

### Removing xenon background

Xenon has several naturally-ocurring isotopes which overlap in mass with the heavier tellurium isotopes (Figure 1a). The argon used in IMC can be contaminated with xenon and as a result some mass channels used to monitor tellurium contain signal from xenon. The amount of xenon in the argon flow varies over time, presumably due to a non-heterogeneous mixture of the gases in the storage vessel. In a phenomenon that is especially disruptive to imaging, large ‘spikes’ of xenon are drawn into the mass cytometer and result in streaks of background signal across a tellurium image (Figure 1b).

**Fig. 1.**
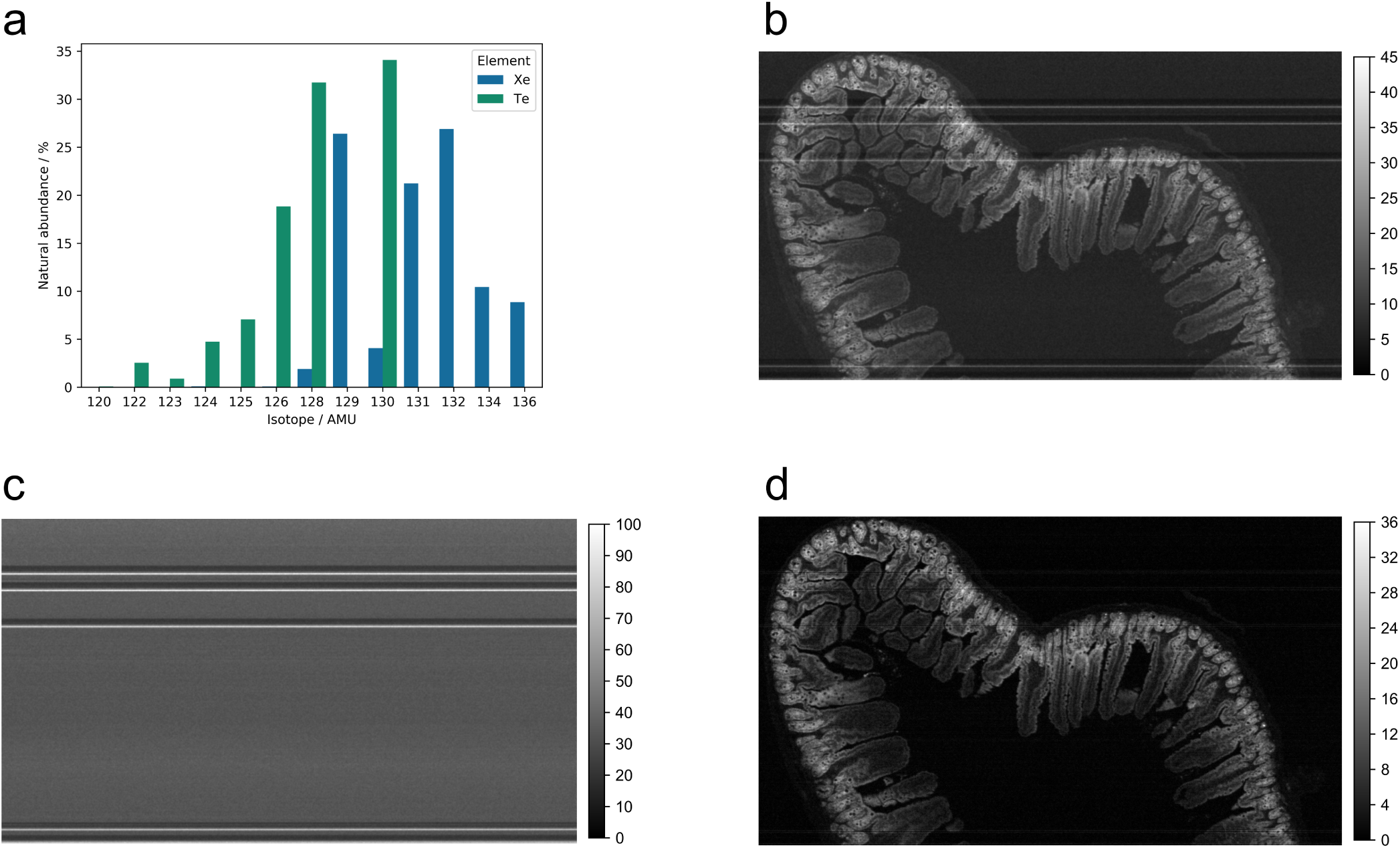
Xenon interference can be removed from tellurium images. (a) Xenon has several isotopes whose masses overlap with tellurium. 128 and 130 are the most obfuscated channels, but ^124^Xe (0.095 % abundant) and ^126^Xe (0.089 % abundant) also contaminate tellurium significantly. (b) The 128 mass channel image, comprising ^128^Te and ^128^Xe. (c) The 134 mass channel image, representing ^134^Xe. This image can be used to estimate the amount of ^128^Xe present. (d) The background subtracted 128 mass channel image, now representing only ^128^Te. Colour bars represent IMC counts in arbitrary units.

To solve this issue, we recorded the 134 AMU image, **X**, which contains xenon but no tellurium, to quantify the xenon in each pixel (Figure 1c). Next we used the known isotopic ratios of ^134^Xe to the xenon contaminating isotope of interest (in this case ^128^Xe) to estimate an image corresponding to the xenon component of the 128 AMU channel, **Y**. By considering that the observed image from the 128 AMU channel, **I**, is a sum of the xenon component **Y** and the tellurium component **T** (i.e. the true ^128^Te image), **T** can be calculated by subtracting **Y** from **I**.

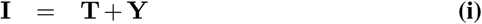

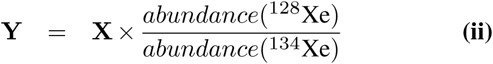

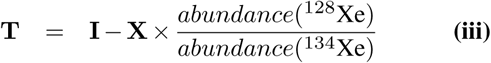

Xenon background subtraction returns an image that represents the corrected tellurium image (Figure 1d). In addition to allowing more accurate visual inspection of images, xenon background subtraction is critical in SLIP experiments where the accurate quantification of each tellurium isotope is important to downstream data analysis.

### Calculating bias in isotope transmission efficiency

Although IMC is a quantitative technique, systematic error exists in the sensitivity of the detector to ions of different mass (9). As a result, the instrument is more sensitive to, for example, ^126^Te ions than ^124^Te ions. While this is of little consequence in natural abundance experiments, SLIP experiments rely on quantifying differences between mass channels and therefore require correction of this transmission efficiency bias (TEB).

We took advantage of data from natural abundance tellurium to calculate the TEB. If the instrument is equally sensitive to all isotopes, the relative intensity of the pixel at location *ij* from the ^*a*^Te image **A** to the pixel at the same location from the ^*b*^Te image **B** would depend only on the relative isotopic abundance:

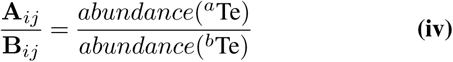

The amount of deviation from Equation iv represents the TEB. We used IMC data from a natural abundance tellurium experiment to calculate the TEB between two isotopes of interest, ^*a*^Te and ^*b*^Te. A ratio image **R** is calculated, in which pixel **R**_*ij*_ is the ratio of the corresponding pixels in the two isotope images of interest, **A**_*ij*_ and **B**_*ij*_:

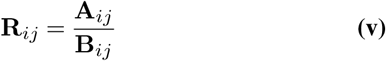

The median of the ratio image **R** is compared to the theoretical ratio of isotope *a* to isotope *b* to give the TEB:

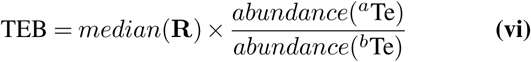

We chose to use a ratio-based approach as opposed to linear regression to avoid the designation of one isotope as the dependent variable and the other as the independent variable. Pixels with a value of 0 in either image must be excluded from analysis to avoid division by 0. Non-zero but low pixel values, while mathematically sound in TEB calculation, skew results by creating unreasonably high or low values in **R**. We found a threshold of 1 count per pixel to allow robust quantification of the TEB. In addition, a moderate Gaussian blur (*σ* = 1 µm) aids in removing outlier pixels from the analysis. As anticipated, removal of xenon background is necessary before calculation of TEB values.

We observed that heavier tellurium isotopes are overdetected by the mass cytometer (Figure 2). This effect has two contributing causes. First, ions with lower mass have lower kinetic energy in the ion beam that travels from the plasma torch to the time-of-flight measurement chamber and have a higher chance of being ejected from the ion beam before detection (10). Second, the mass cytometer is equipped with a quadrupole mass filter that removes ions with *m/z* < 80 Da and is tuned to transmit ions with *m/z* ≈ 160 Da most efficiently (9). Therefore the lower mass Te ions, being further from the target of the mass filter, are transmitted less efficiently.

**Fig. 2.**
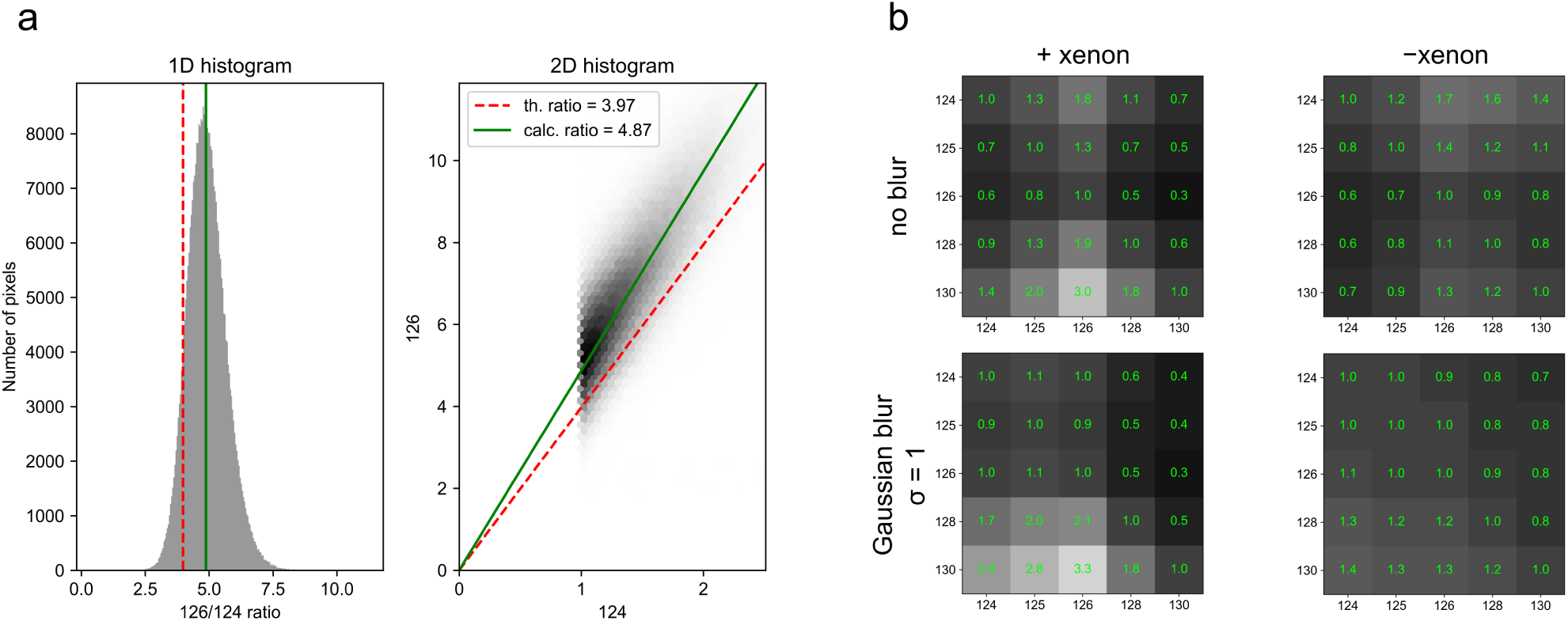
Transmission efficiency bias can be quantified using data from natural abundance tellurium probes. (a) The ^124^Te and ^124^Te channels of the same image are compared to quantify the transmission efficiency bias (TEB). The left panel shows a histogram of the values obtained by dividing the ^126^Te image by the ^124^Te image. The right panel shows the same data as a bivariate histogram. Both panels show the theoretical ratio (based on isotopic abundances) as a dashed line and the calculated ratio as a solid line. (b) Heatmaps displaying the TEB for different pairs of isotopes. TEB values were calculated with or without xenon removal, and with or without Gaussian blurring of the images.

Given that the TEB is a property of the instrument, it is important that users calculate their own values for the TEB between two isotopes, preferably shortly before and after the experiment whose data is to be corrected, since the mass cytometer is known to ‘drift’ over time (11), and individual mass cytometers have different patterns of sensitivity (9).

We applied this approach to data from a published SLIP experiment investigating changes in tumour hypoxia over time (Figure 3) (6). In the experiment, mice bearing PANC-1 xenografts were dosed with two isotopologues of a compound that labels hypoxia 24 hours apart. A difference image is then created, highlighting increasing hypoxia in red and attenuating hypoxia in green. After TEB correction, the overall conclusion of the experiment that cycling hypoxia in tumours occurs without intervention and can be studied with SLIP experiments does not change. However, some regions of the image change colour with TEB correction, indicating a change in the interpretation of their hypoxic shifts. Therefore, TEB correction is an important step in the analysis of data from SLIP experiments.

**Fig. 3.**
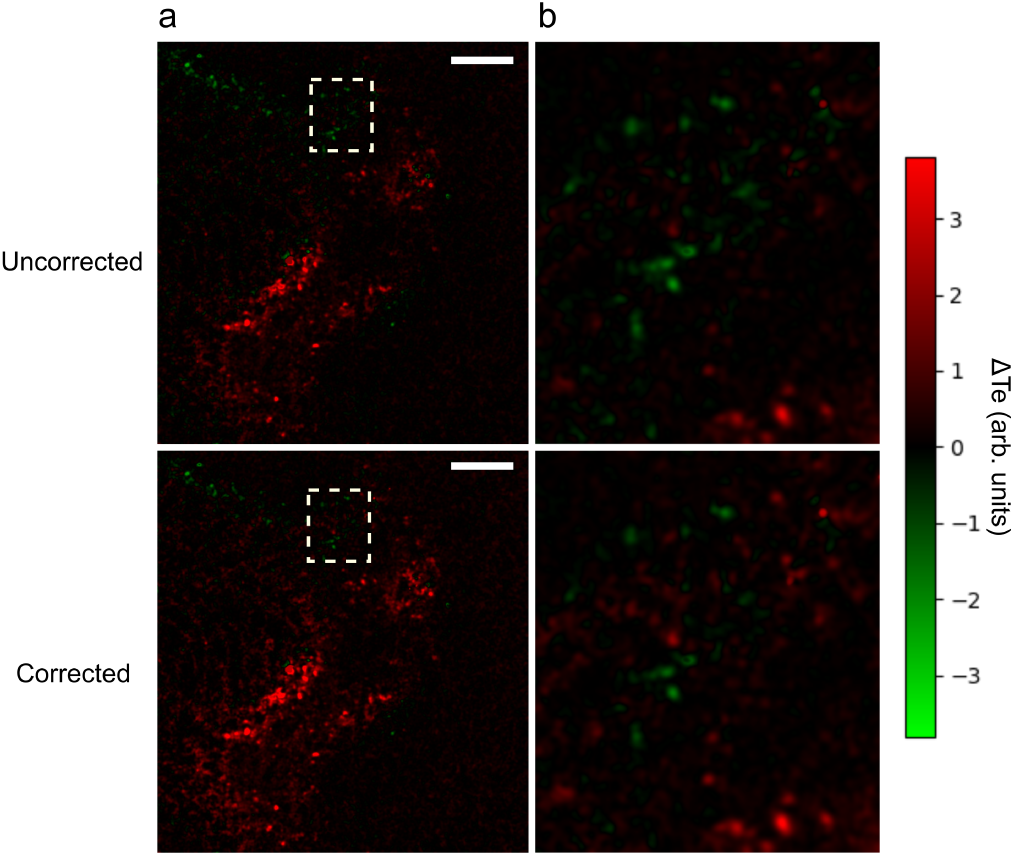
TEB correction is important for accurate SLIP experiments. TEB calculation determined that the instrument is approximately 1.15 more sensitive to ^125^Te than ^122^Te. Using data from a SLIP experiment measuring changes in hypoxia over time, we show the effect of TEB correction on the output image. Column (a) shows the same experiment with or without TEB correction, and the boxed region is expanded in (b). Scale bars are 200 µm.

### Combining tellurium channels for improved signal-to-noise ratio

While SLIP experiments add temporal resolution to IMC, single-timepoint experiments or validation of new probes are carried out with natural abundance tellurium, which has six isotopes with abundances above 1 %. Experiments with natural abundance tellurium may combine images of several tellurium isotopes together to improve signal-to-noise ratio (SNR) and reduce artefacts in the image. There are several possible approaches to combining the tellurium channels. The simplest and most intuitive approach is to sum the channels arithmetically. While this approach is minimally-manipulative, it does not effectively remove background noise. This is because noise in IMC is not distributed around 0, and can only have positive values. Therefore when adding channels together, the noise accrues and does not cancel. An alternative method is taking the product of *n* tellurium channels, then normalizing by taking the *n*^*th*^ root of the product array (i.e. the geometric mean). The geometric approach is more effective at removing background noise from the image because a pixel value of 0 in any channel with result in a pixel value of 0 in the output image. However, if a very low signal image is included in the geometric mean, then the output will be unnecessarily skewed and contain many pixels with a value of 0. Typically, ^130^Te, ^128^Te, and ^126^Te can all be included without skewing the image to 0, since they are the highest abundance isotopes, but users should decide empirically which isotopes to include.

A further option is to stack each image to be combined into a 3-dimensional array and perform a 3-dimensional Gaussian blur. Then the blurred 3-dimensional array can be collapsed into a 2-dimensional image in either an arithmetic or geometric fashion.

We compared the SNRs of an image processed arithmetically against one processed geometrically, both with and without 3-dimensional Gaussian blue (Figure 4). We define SNR as:

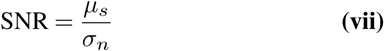

where *µ*_*s*_ is the mean intensity in an arbitrarily-defined signal region of the image, and *σ*_*n*_ is the standard deviation of the intensity in an arbitrarily-defined noise region of the image. The signal region was defined as the region containing tissue mounted on the microscope slide and the noise region as the region containing only the microscope slide.

**Fig. 4.**
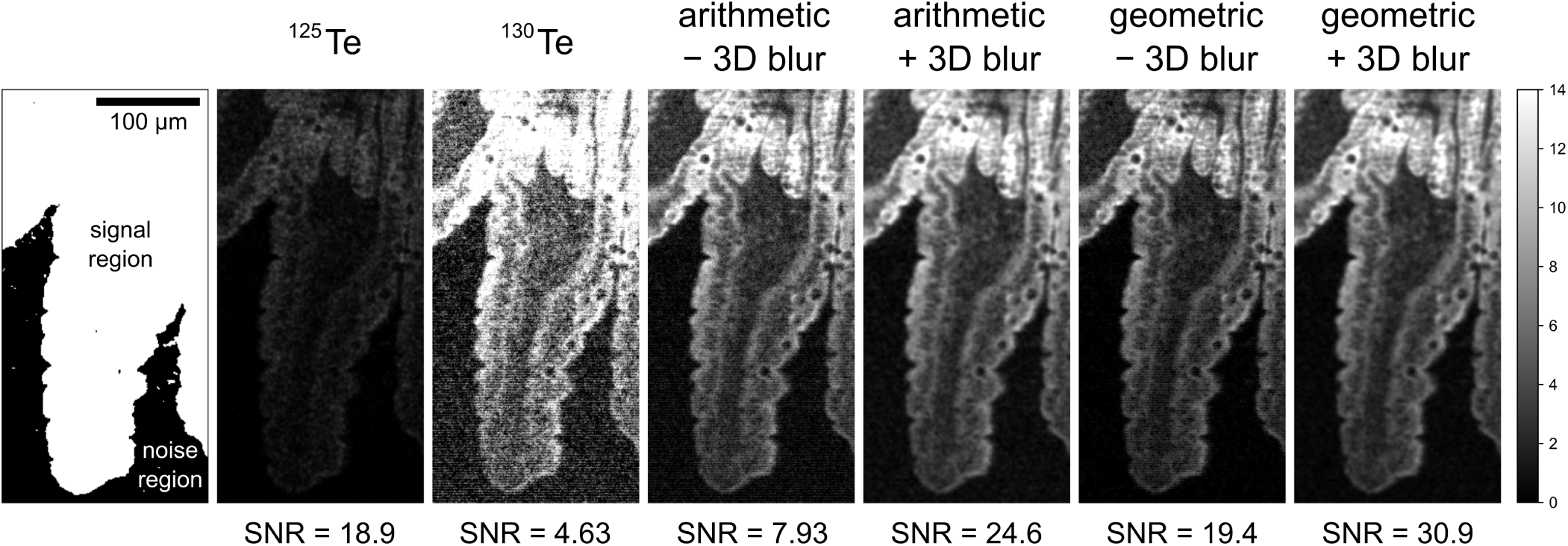
Combination of ^125^Te, ^126^Te, ^128^Te, and ^130^Te images. The signal and noise regions are defined arbitrarily based on knowledge of the tissue of interest. Single isotope images may have either low signal (e.g. ^125^Te) or high noise (e.g. ^130^Te). The effect of combining the images in a arithmetic or geometric fashion, with or without 3-dimensional Gaussian blurring (*σ* = 1) is shown. SNR denotes signal-to-noise ration. Scale is the same in all images and depicted in the left panel.

We found that three-dimensional Gaussian blur provides a strong improvement on the SNR in both arithmetic and geometric modes, but geometric combination improves the SNR more substantially than arithmetic combination. Blurring may improve SNR but will also reduce resolution; we expect that each experiment will have an optimal balance between SNR and resolution that depends on the specific biology being interrogated. Improvement of the SNR with natural abundance probes with this technique will be useful for any experiment involving natural abundance tellurium probes.

## Conclusions

We have described strategies that enhance the analysis of IMC experiments with tellurium probes. We took advantage of the single-mass-unit resolution of IMC and the known isotopic distributions of tellurium and xenon to remove background signal from channels of interest. The polyisotopic nature of tellurium enabled both quantification of instrument biases and enhancement of signal-to-noise ratio of natural abundance tellurium probes.

We expect that the techniques described in this manuscript will become part of the routine IMC workflow when tellurium probes have been used. Coupled with the expanding toolkit of tellurium probes, we hope to progress the study of dynamic biology in vivo.

We have implemented the strategies described here, as well as other general IMC processing methods, in a Python package available at https://github.com/jaybassan/teimc and PyPI (teimc). The package is concise, readable, and amenable to customization; it is open source and based on other open sourced libraries.

We intend to continue the development of software as a complement to laboratory advances, and especially hope to develop cell segmentation solutions coupled with automated SLIP analysis.

## ACKNOWLEDGEMENTS

We thank Fluidigm Inc., the Natural Sciences and Engineering Research Council of Canada, and the Connaught Fund for financial support. We also thank T. Closson for assistance with Imaging Mass Cytometry. We are grateful to N. Law, L. Caporiccio, Dr R. N. Vellanki and Dr B. G. Wouters for preparing the biological samples. This preprint was formatted using a modified class file from the Henriques lab at University College London.

## Bibliography

1. Charlotte Giesen, Hao A.O. Wang, Denis Schapiro, Nevena Zivanovic, Andrea Jacobs, Bodo Hattendorf, Peter J. Schüffler, Daniel Grolimund, Joachim M. Buhmann, Simone Brandt, Zsuzsanna Varga, Peter J. Wild, Detlef Günther, and Bernd Bodenmiller. Highly multiplexed imaging of tumor tissues with subcellular resolution by mass cytometry. Nat. Methods, 11(4):417–422, 2014. ISSN 15487105. doi: 10.1038/nmeth.2869.

2. Qing Chang, Olga I. Ornatsky, Iram Siddiqui, Rita Straus, Vladimir I. Baranov, and David W. Hedley. Biodistribution of cisplatin revealed by imaging mass cytometry identifies extensive collagen binding in tumor and normal tissues. Sci. Rep., 6(November):1–11, 2016. ISSN 20452322. doi: 10.1038/srep36641.

3. Sydney M. Evans, Stephan M. Hahn, Deirdre P. Magarelli, and Cameron J. Koch. Hypoxic Heterogeneity in Human Tumors. Am. J. Clin. Oncol., 24(5):467–472, 2001. ISSN 0277-3732. doi: 10.1097/00000421-200110000-00011.

4. Hanuel Park, Landon J. Edgar, Matthew A. Lumba, Lisa M. Willis, and Mark Nitz. Organotellurium scaffolds for mass cytometry reagent development. Org. Biomol. Chem., 13(25): 7027–7033, 2015. ISSN 14770520. doi: 10.1039/c5ob00593k.

5. Jay Bassan, Lisa M. Willis, Alan Nguyen, Ravi N. Vellanki, Landon J. Edgar, Bradly G. Wouters, and Mark Nitz. TePhe, a tellurium-containing phenylalanine mimic, allows monitoring of protein synthesis in vivo with mass cytometry. Proc. Natl. Acad. Sci. U. S. A., 116 (17): 8155–8160, 2019.

6. Landon J. Edgar, Ravi N. Vellanki, Trevor D. McKee, David Hedley, Bradly G. Wouters, and Mark Nitz. Isotopologous Organotellurium Probes Reveal Dynamic Hypoxia In Vivo with Cellular Resolution. Angew. Chemie - Int. Ed., 55(42):13159–13163, 2016. ISSN 15213773. doi: 10.1002/anie.201607483.

7. Denis Schapiro, Hartland W. Jackson, Swetha Raghuraman, Jana R. Fischer, Vito R.T. Zan-otelli, Daniel Schulz, Charlotte Giesen, Raúl Catena, Zsuzsanna Varga, and Bernd Bodenmiller. HistoCAT: Analysis of cell phenotypes and interactions in multiplex image cytometry data. Nat. Methods, 14(9):873–876, 2017. ISSN 15487105. doi: 10.1038/nmeth.4391.

8. Raúl Catena, Luis M. Montuenga, and Bernd Bodenmiller. Ruthenium counterstaining for imaging mass cytometry. J. Pathol., 244(4):479–484, 2018. ISSN 10969896. doi: 10.1002/path.5049.

9. Sabine Tricot, Mickael Meyrand, Chiara Sammicheli, Jamila Elhmouzi-Younes, Aurélien Corneau, Sylvie Bertholet, Marie Malissen, Roger Le Grand, Sandra Nuti, Hervé Luche, and Antonio Cosma. Evaluating the efficiency of isotope transmission for improved panel design and a comparison of the detection sensitivities of mass cytometer instruments. Cytometry Part A, 87(4):357–368, 2015. doi: 10.1002/cyto.a.22648.

10. “Hongsen Niu and R.S. Houk”. “fundamental aspects of ion extraction in inductively coupled plasma mass spectrometry”. “Spectrochimica Acta Part B: Atomic Spectroscopy”, “51” (“8”):“779–815”, “1996”. ISSN “0584-8547”. doi: “https://doi.org/10.1016/0584-8547(96)01506-6”.

11. Rachel Finck, Erin F. Simonds, Astraea Jager, Smita Krishnaswamy, Karen Sachs, Wendy Fantl, Dana Pe’er, Garry P. Nolan, and Sean C. Bendall. Normalization of mass cytometry data with bead standards. Cytom. Part A, 83 A(5):483–494, 2013. ISSN 15524922. doi: 10.1002/cyto.a.22271.

